# CSAG1 Maintains the Integrity of the Mitotic Centrosome in Cells with Defective P53

**DOI:** 10.1101/778522

**Authors:** Hem Sapkota, Jonathan D. Wren, Gary J. Gorbsky

## Abstract

Centrosomes focus microtubules to promote mitotic spindle bipolarity, a critical requirement for balanced chromosome segregation. Comprehensive understanding of centrosome function and regulation requires a complete inventory of components. While many centrosome components have been identified, others may yet remain undiscovered. We have used a bioinformatics approach, based on “guilt by association” expression to identify novel mitotic components among the large group of predicted human proteins that have yet to be functionally characterized. Here we identify Chondrosarcoma-Associated Gene 1 (CSAG1) in maintaining centrosome integrity during mitosis. Depletion of CSAG1 disrupts centrosomes and leads to multipolar spindles more effectively in cells with compromised p53 function. Thus, CSAG1 may reflect a class of “mitotic addiction” genes whose expression is more essential in transformed cells.

## Introduction

Mitosis accurately and evenly divides the replicated genome into two daughter cells. During mitosis, the spindle forms from microtubules focused by centrosomes at the two opposite poles, and bundles of these microtubules attach to the kinetochores of sister chromatids. Full attachment of microtubules to kinetochores occurs as chromosomes align at the equatorial region in metaphase. Shortly thereafter, at anaphase, the chromatids of each chromosome separate into the two daughter cells [1–3]. Formation and maintenance of the bipolar spindle is critical to ensure proper segregation of chromosomes. The formation of more than two poles during mitosis greatly compromises the fidelity of chromosome segregation at anaphase.

In vertebrate cells, centrosomes are composed of a pair of centrioles surrounded by a condensed cloud of proteins termed the pericentriolar matrix (PCM) [4]. Multipolar mitoses are often caused by centrosome over duplication, failure to cluster extra centrosomes, premature dissociation of centrioles, or PCM fragmentation during mitosis [5]. Some cancer cells with multiple centrosomes can nevertheless form bipolar spindles by clustering the extra centrosomes [6, 7]. In such cells, spontaneous or experimentally induced failure in centrosome clustering leads to multipolarity [6, 8, 9]. The induction of multipolar spindles by microtubule-stabilizing drugs such as Taxol may be a mechanism underlying therapeutic effects in cancer treatment [10]. Overexpression of kinases, polo like kinase 4 (PLK4) and Aurora A, during interphase cause centrosome amplification and multipolarity during subsequent mitosis [11, 12]. Long delays at metaphase induce multipolar spindles in cells that have undergone chromatid separation in cohesion fatigue [13, 14].

In some cases multipolarity in mitosis can occur without centrosome amplification and be caused by centriole disengagement, or fragmentation of the PCM (reviewed in [5]. The presence of damaged DNA may also generate multipolar spindles after cells initiate mitosis with a normal appearing bipolar spindle [15]. Malignant cells with amplified centrosomes often cluster extra centrosomes into two poles to maintain the bipolar spindle and hence avoid massive chromosome instability [7, 16, 17].

Among the many proteins found within the PCM, some have clearly defined functions. For example, γ-tubulin is recruited to the PCM aids in nucleating the microtubules that form the spindle during mitosis [18]. The centrosome becomes enlarged and more defined as the cell approaches prophase of mitosis. How the PCM expands during the preparation for mitosis is poorly understood. The centrosome is not membrane bound and has been suggested to be an example of an organelle formed by phase separation or protein condensation [19–21]. Therefore, there are likely to be components of the centrosome central in maintaining its integrity. Fragmentation of PCM components during mitosis has been reported to occur in a manner that requires the presence of spindle microtubules [22].

In an effort to discover novel mitotic proteins, we used a bioinformatics approach called GAMMA [23]. Briefly, GAMMA processes over 80,000 publicly available high-throughput transcriptional experiments (i.e., microarray and RNA-seq) to identify highly correlated transcripts. Then, using a “guilt by association” approach, even if nothing or little has been published on a mitotic gene, it seems highly likely its transcription would be most strongly correlated with mitotic proteins over other classes. In fact, we and others have successfully used GAMMA to prioritize potentially mitotic proteins for further experimental characterization [24–26].. Using this approach, we identified a mitotic role for a poorly characterized protein, chondrosarcoma-associated gene 1 (CSAG1). mRNA analyses indicate that CSAG1 is highly expressed in chondrosarcoma, other cancers, and in certain normal tissues such as testis and brain (https://www.proteinatlas.org/ENSG00000198930-CSAG1/tissue. Two transcript variants with different 5’ untranslated regions have been described, but both mRNAs code the same 78 amino acid protein [27]. Functions for CSAG1 have not been characterized and, generally, very little is known about regulation of the gene. A closely related gene CSAG2/3, which is also known as Taxol-resistant gene 3, is better studied and shown through RNA analysis to be highly expressed in different cancers [28–30]. Our laboratory has previously shown that depletion of CSAG1 significantly inhibits cell proliferation in a breast cancer stem cell model system [26].

In this study, we functionally characterize the CSAG1 gene product. We report that CSAG1 concentrates at centrosomes and its depletion by siRNA in HeLa cells results in a high level of multipolar anaphase. We found stretched and fragmented PCM in CSAG1-depleted cells suggesting that CSAG1 functions in strengthening the integrity of the PCM during mitosis. In HCT116 cells, we found that cells lacking normal p53 function are more likely to undergo multipolar mitosis when depleted of CSAG1. CSAG1 could be an example of mitotic addiction, reflecting genes whose expression are more indispensable in a p53-compromised background and thus potential targets in cancer therapy.

## Results and Discussion

### CSAG1 depletion results in delayed mitotic progression and multipolar mitotic exit

Live cell imaging of HeLa cells stably expressing GFP-histone H2B revealed that CSAG1-depleted cells initiated mitosis with a bipolar spindle and generally advanced to a normal metaphase. Thereafter, in a large portion of cells, the metaphase plate became bent, indicative of multipolar mitosis, and then cells entered anaphase with the chromosomes segregating into 3-4 distinct DNA masses (Figure 1A and supplemental video 1). Depletion of CSAG1 in HeLa cells caused increased incidence (45%) of multipolar mitotic exit compared to negative control (NC) siRNA (Figure 1B). In addition to the increased frequency of multipolar mitosis, CSAG1 depletion also caused chromosome alignment defects in a small number of cells (Figure 1B). Other mitotic defects such as premature exit from mitosis without alignment or lagging chromosomes were rare. Additionally, CSAG1 depletion increased the elapsed times from nuclear envelope breakdown (NEBD) to anaphase (Figure 1C). Cells transfected with CSAG1 siRNA that did not exhibit the multipolar phenotype also showed delayed NEBD to anaphase compared to control cells (Supplemental figure 1A). The longer duration of progression was attributable to both delays from NEBD to metaphase and from metaphase to anaphase.

**Figure 1:**
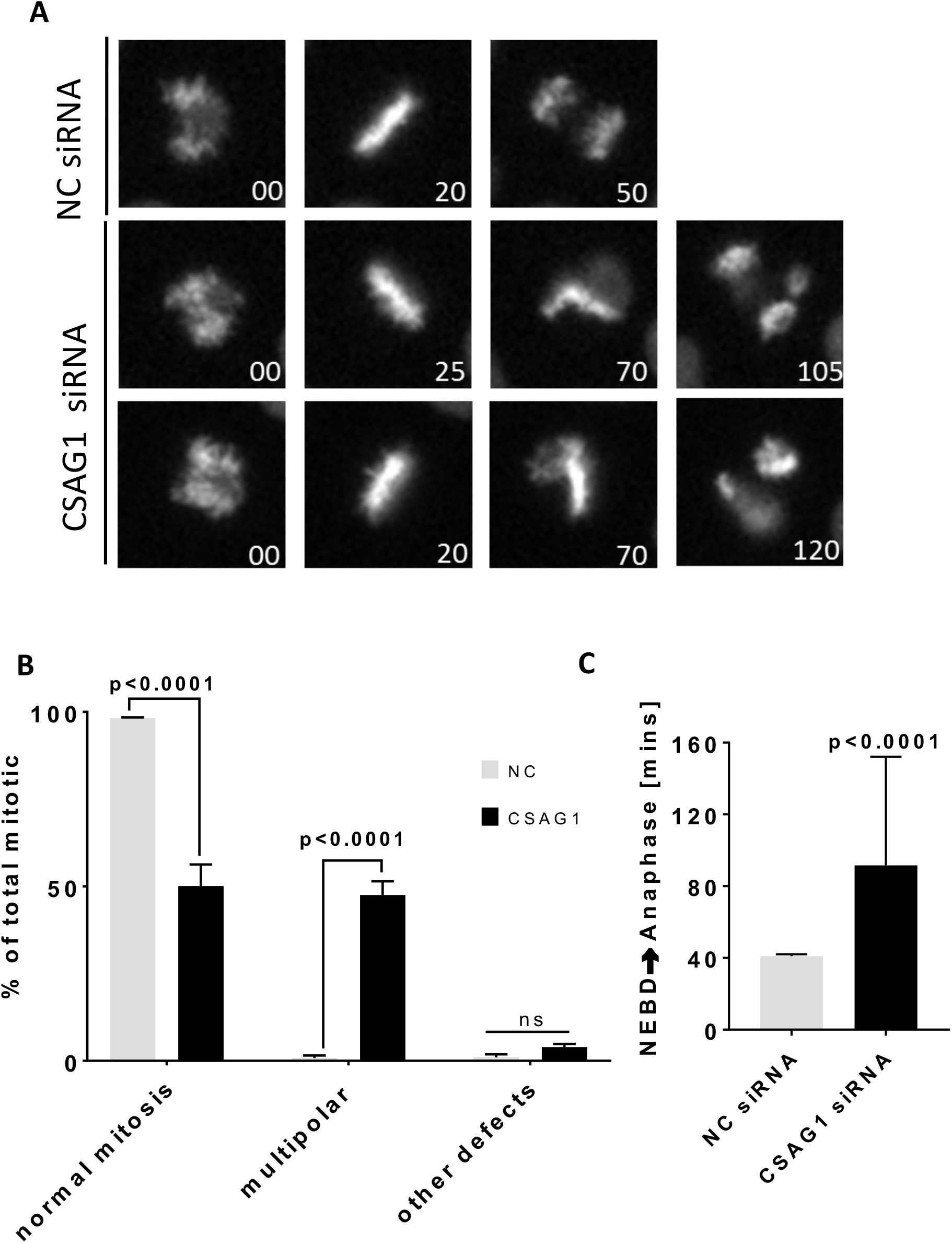
Depletion of CSAG1 by RNAi causes multipolar mitosis. **(A)** Images from live cell microscopy of HeLa-H2B-GFP cells transfected with either negative control siRNA (NC) or CSAG1 siRNA. Top set of panels show an unperturbed mitosis with negative control (NC) siRNA. Bottom two sets of panels show CSAG1 siRNA treated cells exhibiting multipolar mitosis, where the metaphase plate bends shortly after metaphase. **(B)** Cells were analyzed for multipolar spindles of other mitotic defects such as delayed alignment, mitotic exit without alignment, or lagging anaphase chromosomes. Sidak’s multiple comparisons test was used for statistical analysis (**C)** Elapsed times from nuclear envelope breakdown (NEBD) to anaphase were determined in NC or CSAG1 siRNA treated cells. The Mann-Whitney test was used for statistical analysis. Totals of >200 cells from three independent experiments were analyzed. Error bars represent standard deviation CSAG1-depleted cells exhibit multipolar mitosis and take longer to proceed through mitosis.

### CSAG1 accumulates at centrosomes in mitotic cells and expression of siRNA-resistant CSAG1 rescues the multipolar phenotype

To examine localization of CSAG1 we generated a stable cell line with inducible GFP-CSAG1 using HeLa Flp-in TRex cells. In these cells, the region of cDNA insertion is predetermined by the flippase recognition target (FRT) sites, and the amount of protein expression can be regulated by the concentration of added doxycycline [31]. More than 90% of HeLa-GFP-CSAG1 cells showed nuclear localization of the GFP signal in interphase cells when induced with 2 ug/ml doxycycline under live cell imaging conditions (Supplemental figure 2A). In mitotic cells immunofluorescence analysis with anti-GFP antibodies after doxycycline induction showed clear localization of GFP-CSAG1 at spindle poles/centrosomes. Centrosomal localization was confirmed by co-localization with antibodies to pericentrin. Centrosomal localization of CSAG1 peaked at prophase, similar to pericentrin (Figure 2A). These data are consistent with a centrosomal function of CSAG1 in mitotic cells.

**Figure 2:**
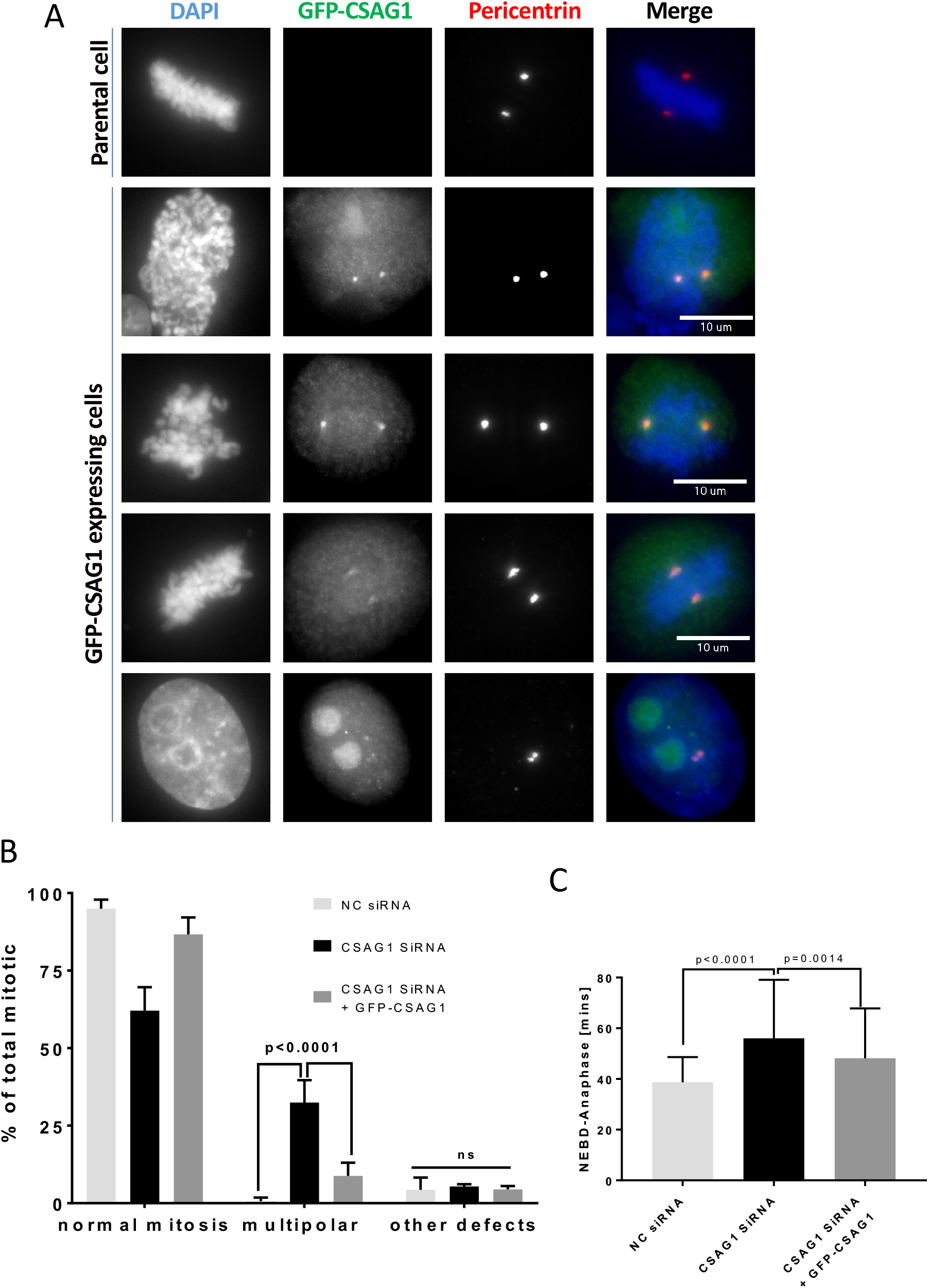
GFP-tagged CSAG1 localizes to spindle pole during mitosis. **(A)** Hela cells stably expressing inducible GFP-CSAG1 were examined for localization of GFP signals during interphase and different stages of mitosis. The top set of panels show absence of GFP signal in uninduced cells. Three other panels show spindle pole/centrosome localization of GFP-CSAG1 during late prophase, prometaphase, metaphase, and interphase respectively. **(B)** The fraction of multipolar mitosis, or other defects were determined in cells from (A) that were transfected with CSAG1 siRNA in presence or absence of doxycycline to induce exogenous siRNA-resistant GFP-CSAG1. Totals of >300 cells were analyzed from three independent experiments. A Two-way ANOVA with Tukey’s multiple comparisons test was used for statistical analysis. **(C)** Elapsed times from NEBD to anaphase was determined in cells from (B). Error bars represent standard deviation, and the Mann-Whitney test was used for statistical analysis.

We used these cells to validate the efficiency of CSAG1 depletion using siRNA targeting the open reading frame of CSAG1 followed by western blotting with anti-CSAG1 antibody. In both soluble lysate and immune-precipitated samples, the GFP-CSAG1 specific band was reduced by >95% after siRNA treatment (Supplemental figure 1C). To exclude non-specific effects of CSAG1 siRNA, we performed rescue experiments. CSAG1 depletion in uninduced GFP-CSAG1 HeLa Flp-in TRex cells caused the multipolar phenotype in approximately 30% of cells. But, when cells were induced with doxycycline to express siRNA resistant CSAG1 and then treated with the smart pool siRNA targeting non-coding regions of endogenous CSAG1, only 6% of cells exhibited multipolar spindles (Figure 2B). Furthermore, mitotic duration was also significantly reduced in GFP-CSAG1 induced cells compared to CSAG1 depleted cells (Figure 2C). These results confirmed that CSAG1 depletion phenotype was indeed due to loss in CSAG1 and not due to non-specific targeting by the siRNA. We also confirmed the siRNA specificity using a different siRNA that targeted the open reading frame (ORF) of CSAG1, which showed similar induction of multipolar spindles but higher levels of chromosome alignment defects, perhaps owing to different levels of depletion (Supplemental figure 1B).

### Increasing mitotic duration and spindle microtubule stabilization exacerbates multipolarity caused by CSAG1 depletion

We noted that CSAG1 depletion in HeLa cells caused a significant delay in mitotic progression even in cells that did not exhibit the multipolar phenotype (Supplemental figure 1A). This observation led us to examine whether delaying mitotic progression experimentally might amplify the multipolar phenotype caused by CSAG1 depletion. We used low concentrations of Nocodazole (25 nM), a microtubule destabilizer, Taxol (1 nM), a microtubule stabilizer, ProTAME (5 uM), an inhibitor of the anaphase promoting complex/cyclosome (APC/C) to delay mitotic progression, and reversine (250 nM), an inhibitor of the MPS1 spindle checkpoint kinase to accelerate it [32]. In control cells, Nocodazole, Taxol, or ProTAME treatment alone increased the time between nuclear envelope break down (NEBD) and anaphase (Supplemental figure 2B) but did not cause multipolar mitosis (Supplemental figure 2C). In contrast, in CSAG1-depleted cells, multipolarity increased to 52% with Nocodazole, 61% with Taxol, and 80% with ProTAME, compared to 45% in control CSAG1-depleted cells (Figure 3A). Treatment of CSAG1 depleted cells with reversine, abrogated the spindle checkpoint, accelerated mitosis, and eliminated the multipolar phenotype. Together these data are consistent with the idea that longer mitotic duration enhances multipolarity induced by CSAG1 depletion.

**Figure 3:**
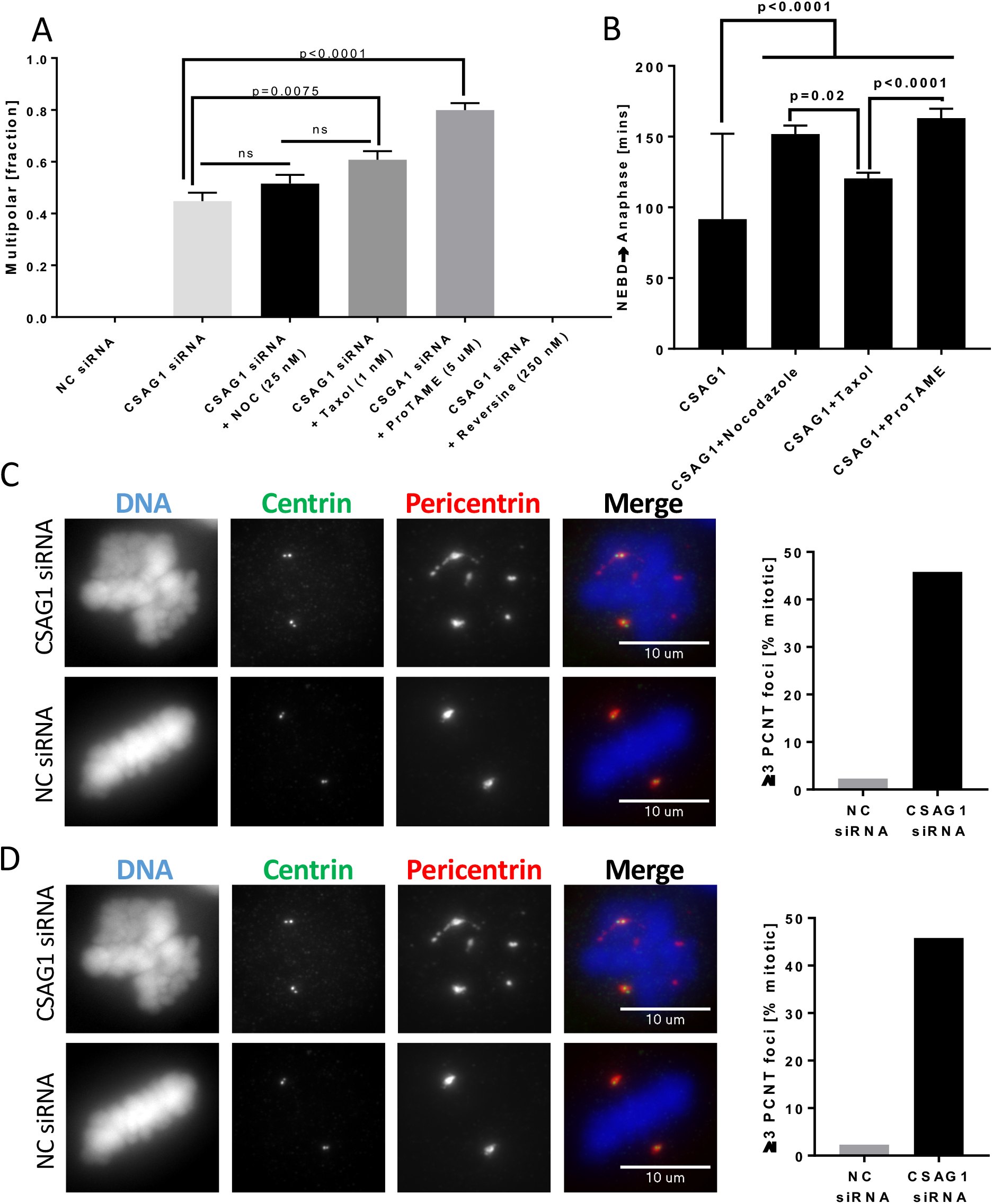
Microtubule perturbation and prolonged mitotic duration increase multipolarity in CSAG1-depleted cells caused by spindle pole fragmentation. **(A)** The fraction of multipolar mitosis was determined in CSAG1-depleted cells treated with low concentrations of nocodazole (5ng/ml), Taxol (1nM), ProTAME (5 uM) or reversine (250 nM). Totals of more than 300 cells were analyzed for each treatment. Error bars represent standard error of mean, Kruskal-Wallis test was used for statistical analysis. **(B)** Elapsed times from NEBD to anaphase was determined in cells from (A). Increasing the duration of mitosis in cells with intact spindle pulling forces increases susceptibility for multipolarity in CSAG1-depleted cells. **(C)** Immunofluorescence images of CSAG1-depleted HeLa cells labeled for γ-tubulin and pericentrin, centrosome markers. The top set of panels shows a CSAG1-depleted cell and the bottom set shows a negative control (NC) siRNA-transfected cell. The graph at right shows the percentage of mitotic cells containing more than 3 γ-tubulin foci. **(D)** Cells as in (A) were immuno-labeled for centrin 1, a centriole marker, and pericentrin. The top set of panels shows a CSAG1-depleted cell and the bottom set shows a control siRNA-treated cell. The graph at right shows fractions of mitotic cells that had more than 3 pericentrin foci. In all cases, centrioles remained paired at two of the spindle poles. For both C and D, more than 100 cells were analyzed. Labeling reveals fragmentation of pericentriolar material induced by CSAG1 depletion.

### Extra spindle poles induced by CSAG1 depletion lack centrioles

Supernumerary centrosomes generate multipolar mitosis which may arise from centrosome reduplication, de-clustering of supernumerary centrosomes, premature centriole separation, or fragmentation of PCM (reviewed in [5]). To determine if CSAG1 depletion induced abnormal centrosome duplication of centrosomes and/or de-clustering of over duplicated centrosomes in HeLa cells, we immunolabeled for γ-tubulin, pericentrin, a PCM component, and centrin 1, a centriole component, in control and CSAG1-depleted HeLa cells. Control HeLa cells exhibited the normal number of centrosomes and centrioles during interphase and mitosis (Supplemental figures 3A and B). Immunofluorescence images also showed that CSAG1-depleted interphase cells had normal numbers of centrosomes and centrioles (Supplemental figures 3A and B). Thus, we found no evidence of preexisting centrosome or centriole over-amplification or CSAG1 depletion-induced over-amplification in HeLa cells. However, CSAG1-depleted mitotic cells with multiple spindle poles showed more than two γ-tubulin foci. These additional pericentrin and γ-tubulin foci lacked centrioles (Figure 3C). In contrast, the supernumerary spindle poles were positive for the both PCM components, γ-tubulin and pericentrin (Figures 3C and D). These data suggest that CSAG1 aids in maintaining PCM integrity during mitosis.

Immunofluorescence data from fixed cell imaging led us to question whether pole fragmentation precedes or follows the bending of metaphase plate detected in our video analysis. We monitored the structure of spindle poles and shape of the metaphase plate during mitosis in CSAG1-depleted cells. We used Hela cells stably expressing GFP–tubulin to reveal spindle poles and applied the far-red DNA dye, sirDNA, for labeling chromosomes. The live imaging revealed that formation of new poles was immediately followed by the bending of the metaphase plate (Supplemental video 1). These findings suggested that bending of the metaphase plate was a consequence of microtubule reorganization in forming multipolar spindles. Taken together these data suggested that CSAG1 depleted cells have normal centrosome number and enter mitosis with a normal bipolar spindle. But extra spindle poles are generated during mitosis at metaphase.

### Loss of CSAG1 alters PCM structure

To test whether CSAG1 depletion disrupts pericentriolar matrix (PCM) which then fragments the poles, we compared distributions of pericentrin in interphase and mitotic HeLa cells after CSAG1 depletion. Interphase cells showed no differences in distribution or the apparent amount of the pericentrin associated with each centrosome (data not shown). However, in mitotic cells, particularly at metaphase, CSAG1 depletion disrupted normal pericentrin organization. Normally pericentrin occupied a well-defined oval shape at the spindle poles in metaphase. However, in some cells this clear definition was not evident and the pericentrin spread to a stretched/non-oval distribution. We classified this type of unstructured and elongated pericentrin labeling as dispersed pericentrin. In control siRNA-transfected cells, 80% of bipolar metaphase cells showed defined, normal appearing pericentrin labeling and only 20% could be classified as dispersed. In contrast, in CSAG1-depleted cells, approximately 70% of bipolar cells showed dispersed pericentrin at poles at metaphase (Figures 4A and B). To better quantify this effect, we measured the axes of PCM shape (as described in figure 4C left) in CSAG1-depleted and control siRNA-transfected cells. In control metaphase cells, the average ratio of length to width of PCM structure was 1.5 while in CSAG1 depleted metaphases the average axis ratio was 2.7. Integrating intensities of immuno-labeled pericentrin showed that the total amount of pericentrin in each pole of CSAG1-depleted cells was comparable to that in control cells (Figure 4D). Overall, CSAG1-depleted cells revealed greater than an 80% increase in axis ratio revealed by labeling with pericentrin antibody (Figure 4C). In extreme cases, CSAG1-depleted cells showed pericentrin labeling up to 5 um from centrioles, something not seen in controls. However, many of these cells retained bipolar spindles and normal appearing metaphase plates. There was no indication of centriole dis-engagement as measured by distance between sister-centrioles (Supplemental figure 3C). We also examined whether changes in PCM distribution/shape in CSAG1-depleted cells might precede the formation of the spindle. However, quantification of PCM axis ratio in prophase cells showed no difference between control and CSAG1 depleted cells (Supplemental figure 3D).

**Figure 4:**
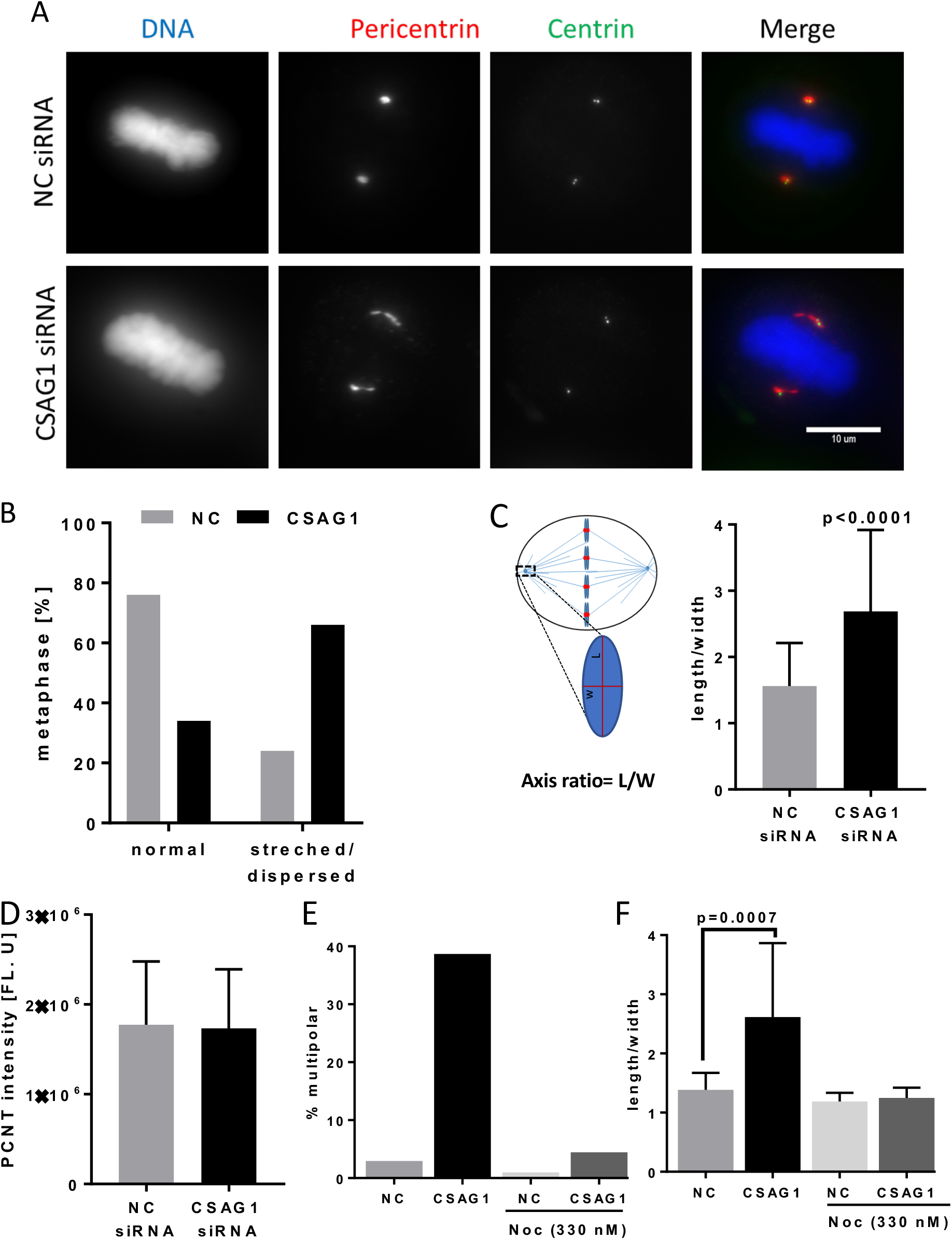
Mitotic spindle microtubules are essential for CSAG1 depletion-induced changes in pericentrin distribution at metaphase. **(A)** Immunofluorescence images of HeLa cells labeled for centrin and pericentrin that were transfected with control siRNA (top set of panels) or CSAG1 siRNA (bottom set of panels). **(B)** The fraction of bipolar metaphase cells with abnormal/dispersed or normal/oval distribution of pericentrin determined in cells from (A). Totals of more than 100 cells were analyzed for each sample. **(C)** Schematic shows method for quantifying of axis ratio (length/width) and the graph shows the ratios calculated. >50 cells were analyzed for each group. **(D)** Total amount of pericentrin was measured from summed z-stack images from B. Multipolarity **(E)** and pericentrin shape/distribution **(F)** was evaluated in CSAG1-depleted cells with or without Nocodazole. CSAG1 depletion causes abnormal pericentrin shape/distribution in bipolar metaphase cells, does not alter the amount of pericentrin at poles. The redistribution requires intact microtubules. The Mann-Whitney test was used for statistical analysis and error bars show standard deviation.

The timing of the altered PCM shape/distribution coincided with formation of mitotic spindles in prometaphase/metaphase. We examined whether spindle microtubules were required for pole fragmentation and/or PCM disturbance. To test this, we disrupted microtubules with a high concentration of Nocodazole (330 nM) for 6 hr in control and CSAG1-depleted cells. This treatment abolished the formation of multiple poles and the changes in PCM shape and distribution (Figure 4E and F). Thus, PCM fragmentation induced by CSAG1 depletion is dependent on the presence of spindle microtubules.

To test the possibility that PCM shape change and fragmentation was caused merely by metaphase delay, we examined the PCM distribution pattern in cells arrested at metaphase with MG132-treatment and found no change (Supplemental figure 3E). These data suggest that CSAG1 depletion compromises PCM and causes it to become dispersed along the microtubules at the poles rendering it susceptible to fragmentation and formation of acentriolar supernumerary poles, which generate multipolar mitotic spindles at high frequency.

### The multipolar phenotype is enhanced by compromised p53 function/expression

A study of 56 different human cell lines shows large variations in CSAG1 transcript levels [33]. In some cell types CSAG1 transcript levels are undetectable while in others, transcript levels are up to 80 reads per kilo base (RPKB) [33].This finding suggested that CSAG1 might not be essential for mitosis in all cells but might serve important roles in certain types of transformed cells. In particular, the RNAseq data revealed that CSAG1 transcript levels were substantially higher in malignant melanoma cell lines.

To examine whether CSAG1 is an essential protein for mitotic progression in non-transformed cells, we depleted CSAG1 in RPE1 cells which are immortalized through expression of telomerase but are otherwise considered to reflect normal, non-transformed cells [34, 35]. RPE1 cells depleted of CSAG1 did not show mitotic defects, though mitotic progression was marginally delayed (Supplemental figure 4A and B), and a small fraction of CSAG1 depleted RPE1 cells exited mitosis without complete chromosome alignment (Supplemental figure 4A). As a control for transfection efficiency, RPE1 cells were depleted of the essential mitotic regulator, Polo like kinase 1 (PLK1), required for bipolar spindle formation. Nearly every PLK1 depleted cell arrested in mitosis due to spindle collapse (Supplemental figure 4A).

Among many differences between RPE1 and HeLa cells is their p53 status. RPE1 cells have normal p53 function whereas in HeLa cells p53 is degraded via the papilloma virus E6 protein. We hypothesized that p53 function might be critical for unmasking the CSAG1 depletion phenotype. For these experiments, we depleted CSAG1 in HCT116 ‘wild type’ cells, which express p53, and in a subline where both copies of the p53 gene were inactivated (HCT116 KO) [36]. Similar to effects with HeLa cells, CSAG1 depletion in HCT116 KO cells caused multipolar mitotic exit in about 40% of mitoses. In contrast, only 5% of HCT116 parental cells showed multipolar mitoses (Figure 5A). Mitotic progression was delayed in both the parental and p53 KO cells, but the delay in the p53 KO cells was more severe. HCT116 parental cells were delayed by about 8 mins, but the p53 KO cells were delayed by 22 mins (Figure 5B).

**Figure 5:**
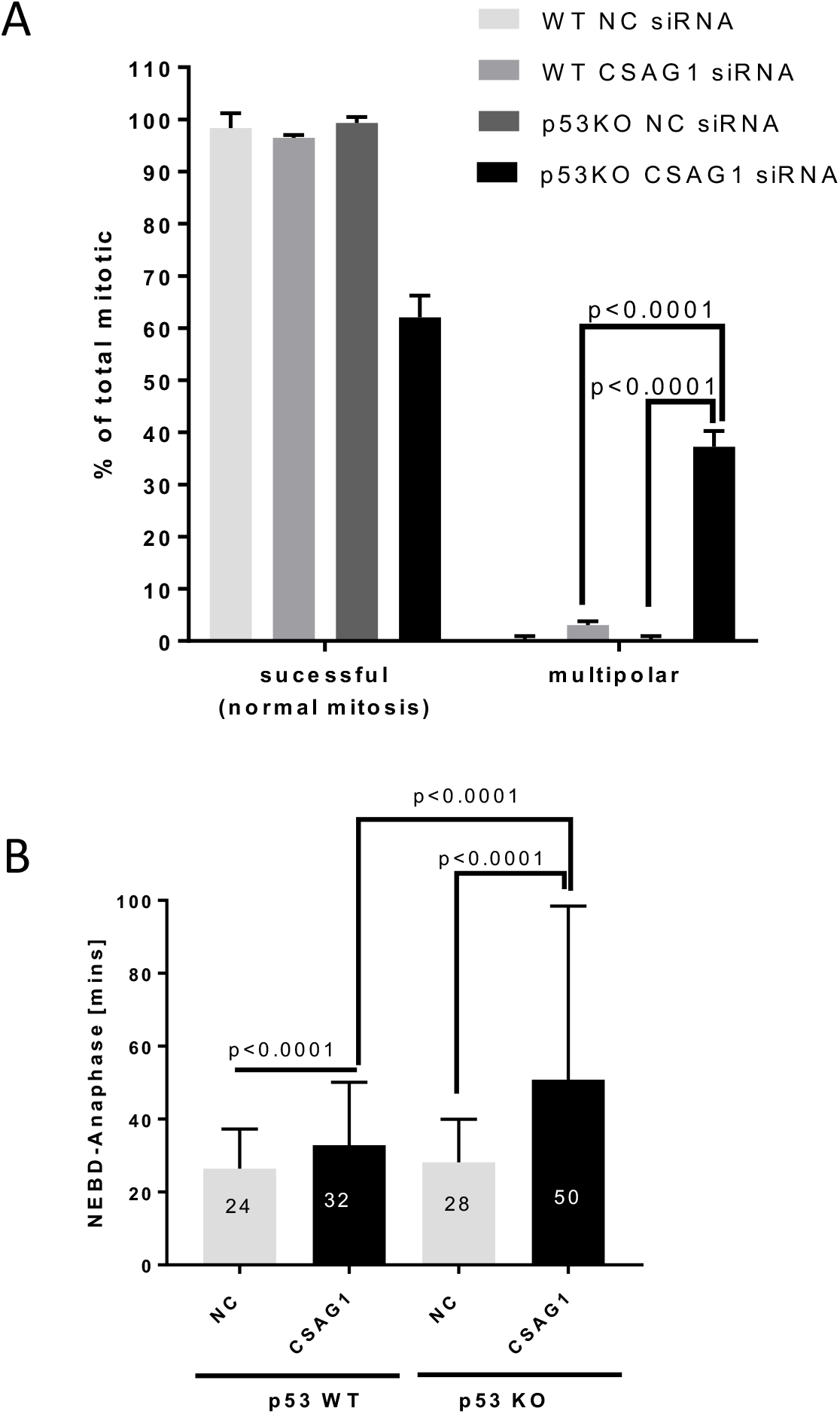
Multipolar mitosis due to CSAG1 depletion is affected by p53 expression. **(A)** The percentage of multipolar mitotic cells was determined in HCT116 parental or p53 KO cells after treatment with NC or CSAG1 RNAi. Totals of >300 cells were analyzed for mitotic defects from three independent experiments. Error bars represent standard deviation. Two way ANOVA with Sidak’s multiple comparisons test was used for statistical analysis. **(B)** Elapsed times from NEBD to anaphase were determined for cells from (A) that did not show mitotic defects. Totals of >200 cells from three independent experiments were analyzed. CSAG1 depletion caused pronounced multipolar mitosis in p53 KO cells and increased mitotic duration in both HCT 116 parental and p53 KO cells.

We have characterized mitotic functions for a previously uncharacterized gene, CSAG1. Exogenously expressed CSAG1 localizes to the centrosome throughout the cell cycle becoming more concentrated there in mitosis. Using siRNA depletion coupled with live cell imaging, we show that CSAG1 plays an important role in maintaining mitotic spindle pole integrity. Overall, our study establishes CSAG1 as a centrosomal protein that maintains pericentriolar integrity, particularly in p53-deficient cells. Unfortunately, we were unsuccessful in detecting endogenous CSAG1 protein because both an antibody we prepared to bacterially expressed CSAG1 and a commercial antibody failed to generate positive signals by Western blotting or immunofluorescence. However a study examining post-mortem brain tissue had detected a phospho-peptide specific to CSAG1 protein via mass-spectrometry [37]. Combined these findings suggest that the endogenous protein may be expressed at very low levels.

Multipolarity in mitosis leads to massive missegregation. Multipolarity has several potential sources, e.g. centrosome over amplification, failure to clusters extra centrosomes, and premature centriole disengagement. None of these were evident in CSAG1-depleted HeLa cells. CSAG1-depleted cells enter mitosis with two normal spindle poles, but these often exhibit extended areas of PCM compared to control cells. Immunofluorescence data show co-localization of pericentrin and GFP-CSAG1 (Figure 2). With time in many CSAG1-depleted cells, supernumerary poles arise that lack centrioles but contain PCM components including pericentrin and γ-tubulin. Thus, the new poles form by PCM fragmentation from the original spindle poles (Figures 3C and D).

Although, many proteins cluster at centrosomes, it remains unclear how the integrity of the centrosome is maintained, particularly in light of the mechanical forces imposed during chromosome segregation in mitosis. Previously, the protein Kizuna was also shown to be essential for maintaining PCM integrity. Recent ideas suggest that centrosomes assemble through phase separation, which may be cell cycle regulated [19–21]. One study suggests that protein phosphatase 4c promotes centrosome maturation and microtubule nucleation [38]. Centrosome maturation and PCM expansion during mitosis is also believed to occur via phase separation [20, 21]. We propose that CSAG1 contributes to mitotic centrosome PCM integrity perhaps through a role in phase separation. In the absence of CSAG1 forces dependent on spindle microtubules at the pole cause disruption and formation of supernumerary poles. The exact molecular mechanisms through which it stabilizes PCM are subjects of current and future inquiry. While multipolarity was enhanced by treatments that delayed anaphase onset in CSAG1-depleted cells, arrest at metaphase by itself does not induce spreading of PCM at poles. HeLa cells delayed at metaphase with the proteasome inhibitor, MG132 did not show altered pericentrin distribution, indicating that centrosomes do not spontaneously fragment to form multiple poles after short mitotic delays (Supplemental figure 3). An additional effect detected with CSAG1 siRNA treatment, particularly evident in RPE1 cells and HCT116 cells was repression of entry into mitosis by 60 to 70%. Because unlike multipolarity, this effect was not rescued by expression of wild type CSAG1, we believe it likely reflects an off-target effect of our siRNA treatment. siRNA targeting the ORF does not reduce the mitotic entry, further strengthening the notion that mitotic entry phenotype is indeed a side effect of particular siRNAs targeting the UTR of CSAG1.

Evidence from cells expressing normal p53 suggest that CSAG1 is not strictly essential in normal cells though it may increase the fidelity of chromosome segregation. Even in HeLa cells, which lack p53 function, only 45% of CSAG1-depleted mitotic cells show multipolarity. One explanation is that CSAG1 may be more critical in malignant cells that often exhibit mitotic delays. Some transformed cells take longer to proceed through mitosis, and those cells are more likely to exhibit multipolarity when centrosome structure is compromised. The precise role that p53 plays in this process is uncertain though other work has shown a correlation between p53 inactivation and sensitivity to depletion of mitotic regulators [39].

At first glance, multipolarity in CSAG1-depleted cells might be attributed to delays in mitotic progression. However, extended mitotic duration does not fully explain the extent of multipolarity observed with CSAG1 depletion. Cells treated with low concentrations of Nocodazole, Taxol, or ProTAME exhibit mitotic delay equal to or longer than those induced by CSAG1 depletion but show no/low levels of multipolarity. Instead we attribute the increased multipolarity in CSAG1-depleted cells to weakened PCM integrity which fragments during mitotic delays. Different means of mitotic delays differently affect the frequency of multipolarity depending upon the spindle force exerted on PCM. In CSAG1 depleted cells, Nocodazole (25 nM) treatments slows mitosis more than Taxol (1nM) treatment but nocodazole treatment causes less multipolarity than Taxol treatment. Nocodazole (25 nM) treatment, which should partially destabilize microtubules and weaken the spindle pulling forces did not significantly increase the frequency of multipolar mitosis in CSAG1-depleted cells. While, Taxol (1 nM) treatment, which stabilizes microtubules and potentially strengthen the spindle forces caused more multipolarity with much shorter mitotic delays. The critical role of spindle microtubules in PCM fragmentation and multipolar mitosis is further illustrated by ProTAME (5 uM) treatment. ProTAME treatment delayed cells at metaphase and doubled the frequency of multipolarity compared to CSAG1 depletion alone (Figure 3A). De-polymerization of spindle microtubules abolishes the multipolarity in CSAG1 depleted cells indicating that dynamic microtubules are necessary for pole fragmentation (Figure 4F).

The connection between the multipolar phenotype and p53 is interesting from both a mechanistic level and as a lead in identifying cancer-specific targets. No correlation between p53 and CSAG1 expression is detected suggesting that one gene is not regulating the other gene at the transcriptional level [33]. Cells depleted of centrosomal components such as pericentrin, γ-tubulin and cNap1 appear to arrest at G1 by a p53-mediated G1 checkpoint [40]. This explanation perhaps explains why CSAG1-depleted HCT116 cells and RPE1 cells with normal p53 may exhibit a subtle flaw in centrosome structure that causes arrest or delay in interphase, resulting in reduced mitotic entry. However, p53 knock-out HCT116 cells also showed reduced entry into mitosis. Potentially a distinct p53-independent interphase checkpoint pathway may also function in this situation to detect centrosome abnormalities.

Overall, our data establish CSAG1 as a centrosomal protein which aids in maintaining spindle pole integrity. Defects are notably seen in transformed p53-deficient cells and enhanced by delayed mitotic progression. Identification of interacting partners and specific molecular functions of CSAG1 will provide better understanding of how spindle pole integrity is maintained.

## Materials and Methods

### Cell Culture

HeLa, RPE1 and HCT116 cells were cultured in flasks in DMEM-based media supplemented with 20 mM HEPES, non-essential amino acids (NEAA), sodium-pyruvate, 1X penicillin-streptomycin (P/S, Corning, 30-002-CI) and 10% FBS. Cells were maintained at 37°C in 51% CO_2_ in a water-jacketed incubator. Cells were subcultured by trypsinization every other day. Mycoplasma contamination was routinely tested cytologically. All cell lines used in this study were mycoplasma free.

### Live cell imaging and drug treatment

Cells grown in 4 or 8 well cover glass chambers were transfected with 100 nM of either negative control siRNA (cat# D001810-10-20) or CSAG1 siRNA pool from Dharmacon. 30-36 h post transfection, media was replaced with Fluorobright fluorescent imaging media containing 500 nM sirDNA (Cytoskeleton). The chamber was then transferred to a Nikon Ti microscope fitted with an OKOlab environmental control chamber. Images were acquired every 7-10 mins using a Nikon 20X objective and Nikon CMOS camera. After completion of imaging, files were exported as TIFF files and analyzed with Metamorph software. Elapsed times from nuclear envelope breakdown (NEBD) to metaphase and to anaphase were determined for each cell that entered mitosis. Additionally, each cell entering mitosis was examined for mitotic defects such as chromosome misalignment, mitotic exit without alignment, lagging chromosomes, anaphase bridges, multipolar mitotic exit, and formation of micronuclei.

For drug treatment, HeLa cells transfected with control or CSAG1 siRNA were treated with either 25 nM Nocodazole, 1 nM Taxol, Reversine (250 nM) or 5 uM ProTAME at beginning of live cell imaging.

### Immunofluorescence

HeLa cells were grown in 22 mm^2^ coverslips in 6 well plates. 24 h after seeding, cells were transfected with 50-100 nM of CSAG1 siRNA. 36-48 h post siRNA transfection, cells were fixed with either −20°C methanol at −20°C for 20 mins or co-fixed/extracted with 2% PFA + 1% TX-100 in 1X PHEM (60 mM <u>P</u>IPES, 25 mM <u>H</u>EPES, 10 mM <u>E</u>GTA and 4 mM <u>M</u>gCl_2_) buffer for 15 mins. Cells were then blocked with 20% boiled normal goat serum (BNGS) for 30 mins then incubated with primary antibodies. Rabbit anti-pericentrin antibody (cat# Ab4448, Abcam) was used at 1:2500, mouse anti-Y-tubulin (cat# T5326, Sigma) was at 1:300, mouse anti-centrin 1 (cat# 04-1624, EMD Millipore) at 1:2500 and anti-GFP Nano-body was used at 1:300 (Chromotek). Cells were washed 3 times for 5 mins with MBST (MOPS buffered saline with 0.05% Tween 20), then incubated with secondary antibodies; FITC conjugated to Goat anti-Rabbit IgG (1:800), Cy3 conjugated to Goat-anti-Mouse IgG (1:1500) for 2h at room temperature. Cells were washed twice again then stained with 200 ng/ml of DAPI for 3 mins. They were then washed and mounted in Vectashield, and coverslips were sealed with nail polish. Images were acquired using a Zeiss Axioplan II microscope with either 63X or 100X objectives, Hamamatsu ORCA II camera, and Metamorph software.

### Generation of GFP-CSAG1 cell lines

The CSAG1 coding region was amplified from a Gateway entry vector plasmid obtained from DNASU to which a stop codon was added. The insert was cloned to a Topo D entry vector, then fused to a MYC-GFP-plasmid with pcDNA5/FRT/TO destination vector modified for gateway cloning. HeLa Flp-in TRex cells were transfected with 900 ng of flippase (pOGG44) plasmid and 100 ng of GFP-CSAG1 plasmid in a 6 well plate. After 48 h, cells were transferred to a 15 cm plate then selected with 200 ug/ml Hygromycin for 20 days. Surviving clones were pooled, treated with doxycycline to induce CSAG1 for 48 h then FACS sorted by fluorescence for GFP expression using a FacsAriaIIIu. These cells were maintained in Hygromycin selection throughout the study.

For rescue experiments, GFP-CSAG1-HeLa Flp-in TRex cells were grown in chambered coverslips for 24 h with or without 2 ug/ml of doxy then transfected with either control siRNA or CSAG1 siRNA. 30-36 h later cells were imaged as described earlier, and quantified.

For CSAG1 localization, GFP-CSAG1-HeLa Flp-in TRex cells were grown in coverslips with or without doxycycline (2 ug/ml) for 48 hours. Then, cells were co-fix extracted with 2% PFA +1% TritonX-100 in 1X PHEM buffer for 15 minutes at room temperature. Fixed cells were washed once then labelled with anti-pericentrin antibody and GFP binding protein (nano-body) fused to Atto 488 (1:300). The protocol described earlier was followed for slide preparation and image acquisition.

### Short Metaphase arrest

Hela cells were treated with 20 uM MG132 for 1.5 h then fixed with −20°C methanol as described earlier. Then cells were labelled with anti-pericentrin antibody and then with DAPI. Metaphase cell were randomly selected from untreated control cells and MG132 treated cells then imaged. Pericentrin distribution patterns were scored manually.

### Antibody production and validation

GST-tev-CSAG1 fusion protein was produced in a bacterial expression system, purified, and then sent to Cocalico for immunization. Serum was tested periodically by western blot for GST and CSAG1 protein. After confirming the presence of GST and CSAG1 specific antibodies in the serum, the serum was collected then affinity purified using His-CSAG1 linked to the amino link plus coupling resins (Thermo scientific, #20501) according to manufacturer’s protocol. Serum was tested again by blotting CSAG1 recombinant protein for specificity and purity of purified antibodies. Serum specificity/reactivity was also tested in immune-precipitated GFP-CSAG1 cell lysate. Clearfield cell lysate was IPed using GFP-Trap-A beads from Chromotek then the blotted with purified serum. Uninduced samples (no doxycycline treatment) of HeLa-GFP-CSAG1 cells were negative controls.

### Immunoprecipitation

For IP using GFP-Trap-A beads (ChromoTek, # GTA-20), the manufacturer’s protocol was followed with slight modification. Briefly, inducible GFP-CSAG1 HeLa cells were grown in 10 cm tissue culture dishes for 48 h with or without 2ug/ml of doxycycline then cells were harvested by trypsinization, washed in cold PBS then lysed in IP buffer (10 mM Tris-HCl pH 7.4, 150 mM NaCl, 1 mM EDTA, 0.5% NP-40 and 1:200 Protease inhibitor (Sigma, #P8340)). Cells were lysed by vigorous pipetting every 10 minutes for total of 30 mins then centrifuged at 16000 Xg for 12 mins. Clear supernatant was transferred to a new micro centrifuge tube then 25 ul of pre-equilibrated anti-GFP beads were added to the sups and incubated with mixing for 2 h at 4C. Then, the beads were centrifuged at 200Xg for 2 mins and supernatant was discarded. The beads were washed 3X with 500 ul of IP buffer, then 30 ul of 3X LDS loading buffer with 150 mM DTT was added directly to the beads, and samples were used for SDS-PAGE.

For western blots, 1X loading sample was prepared using 4X LDS (Thermofisher scientific, #NP008) sample buffer + 100 mM DTT (final). The cell lysates (whole cell lysate, supernatant, or IPed samples) were treated at 95°C for 10 mins. The samples were loaded in 4-12% Bis-Tris NuPage SDS gel and electrophoresed in MOPS SDS buffer at 120V until the dye front reached the bottom of the gel. Samples were transferred to 45 micron immobilon-FLPVDF membrane (Millipore, # IPFL00010), blocked with 1% fish gelatin in PBST(0.5% Tween-20) for 30 mins, then incubated with in house anti-CSAG1 antibody (1:100) overnight at RT and washed 3 times with PBST and incubated with far red goat anti-rabbit IgG IR700 (1:10000) from (Azure,AC2128).

## Acknowledgement

We thank Dr. Andrew Holland for HeLa-flip-In TRex cells. We would also like to thank all the members of Program in Cell Cycle and Cancer Biology at Oklahoma Medical Research Foundation (OMRF) for insightful discussion and comments. This study was supported by grants R35GM126980 (GJG) and P20GM103636 (JDW) from the National Institute of General Medical Sciences and by the McCasland Foundation.

---

**Supplemental figure 1:**
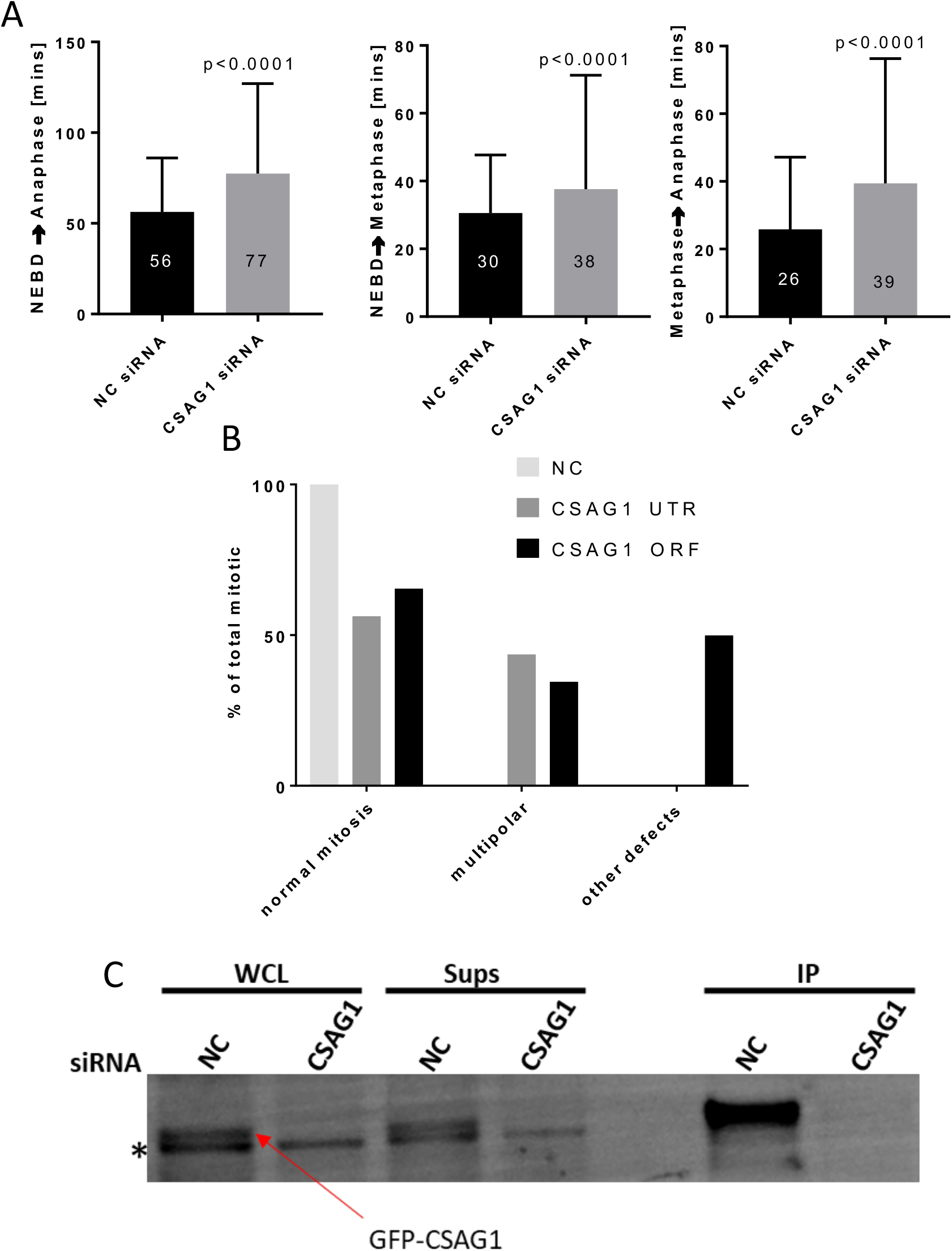
CSAG1 depletion causes mitotic delay in cells that do not show multipolar phenotype. **(A)** Elapsed times from NEBD-anaphase (left), NEBD-metaphase (middle), and metaphase-anaphase (right) were determined in CSAG1-depleted HeLa-H2B-GFP cells. The graphs were plotted excluding the cells that showed multipolar mitosis. Totals of >180 cells from three independent experiments were analyzed. The Mann-Whitney test was used for statistical analysis. **(B)** Mitotic multipolarity or other mitotic defects were assessed in HeLa cells transfected with negative control siRNA (NC) siRNA targeting the UTR of CSAG1 or siRNA targeting an ORF region of CSAG1. Totals of >100 cells were analyzed per siRNA transfection. **(C)** Western blot of whole cells lysate (WCL), soluble fraction (sups) and immuno-precipitated samples (IP) of HeLa cells expressing GFP-CSAG1 that were transfected with either control siRNA or CSAG1 siRNA targeting the ORF. GFP-CSAG1 in significantly enriched after IP in control but not detected in CSAG1 siRNA samples. The same band is completely removed with CSAG1 siRNA in WCL and sups samples. * shows a nonspecific band that was not altered with CSAG1 siRNA.

**Supplemental figure 2:**
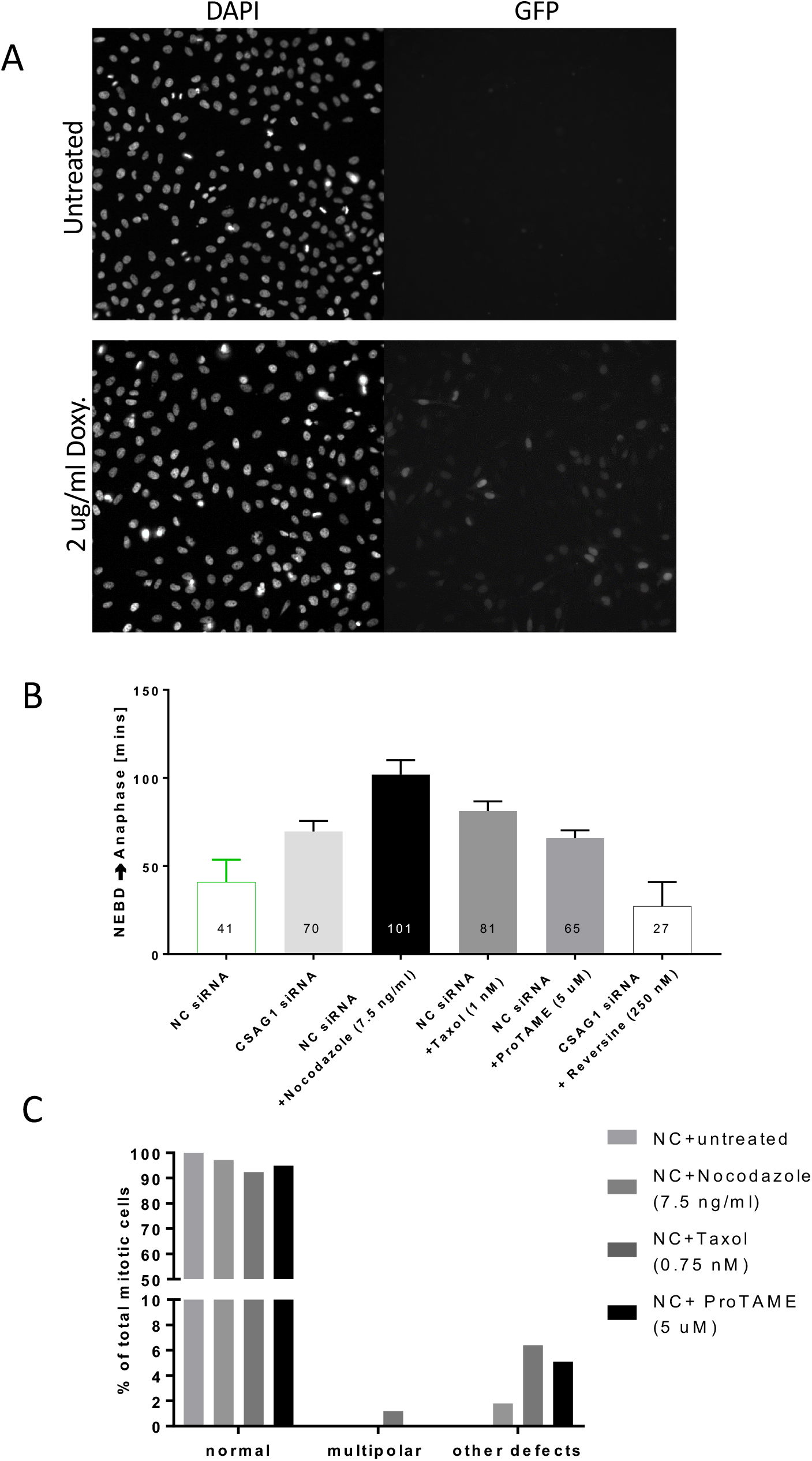
Subtle modulations of mitotic duration with low concentrations of drugs do not induce significant levels of multipolar spindles or other mitotic defects in control cells. **(A)** HeLa cells expressing inducible GFP-CSAG1 were analyzed by live cell imaging for 24 h with or without pretreatment with 2 ug/ml doxycycline. The upper panels show untreated cells and lower panels show doxycycline-treated cells. About 90% of cells were GFP positive after doxycycline treatment whereas 0% were positive in the absence of doxycycline. **(B)** Elapsed times from NEBD to anaphase **(C)** multipolar mitosis were determined in HeLa cells that were transfected with NC siRNA and treated with either nocodazole (25 nM), Taxol (1 nM), ProTAME (5 uM) or reversine (250 nM). Totals of >200 cells for each treatment were analyzed. Numbers within each bar show the average times from NEBD to anaphase.

**Supplemental figure 3:**
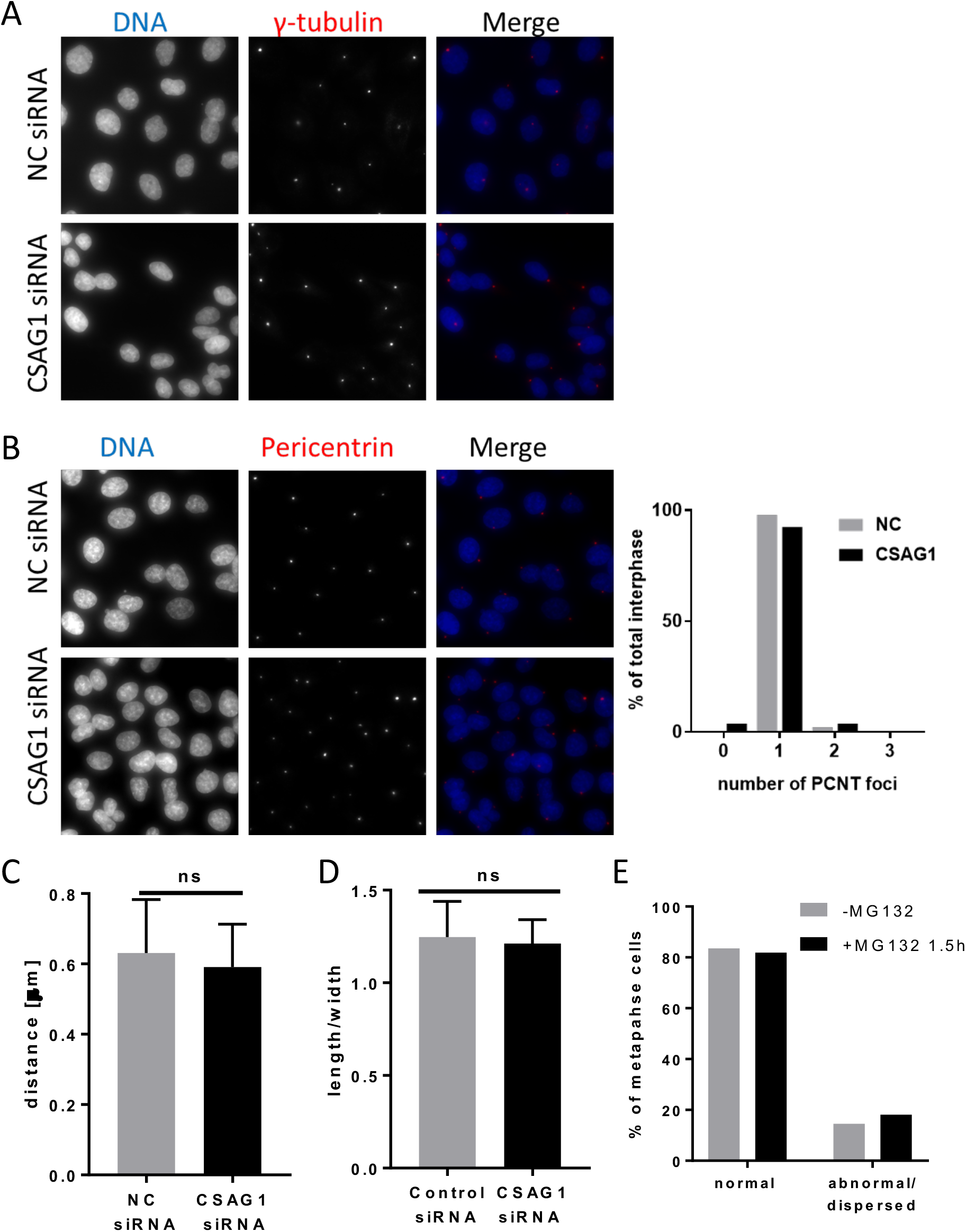
Interphase centrosome numbers and centriole separation are not altered in CSAG1-depleted cells, and short delay at metaphase does not change pericentrin distribution at spindle poles of control cells at metaphase. The total number of centrosomes in interphase cells was determined using **(A)** γ-tubulin or **(B)** pericentrin labeling of HeLa cells. The graph at right shows the frequency distribution of pericentrin foci in interphase cells. Totals of >100 interphase cells were examined for each sample. CSAG1 depletion does not cause increased numbers of centrosomes in interphase. (C) Distances between centrioles were measured in spindle poles of metaphase cells transfected with negative control (NC) or CSAG1 siRNA. **(D)** PCM axis ratios were measured in early prophase cells depleted of CSAG1. **(E)** Pericentrin labeling was evaluated as normal or abnormal/dispersed in metaphase HeLa cells treated with MG132 for 1.5 h. Totals of 150 cells were examined for each sample. Metaphase delay induced by MG132 does not alter pericentrin distribution.

**Supplemental figure 4:**
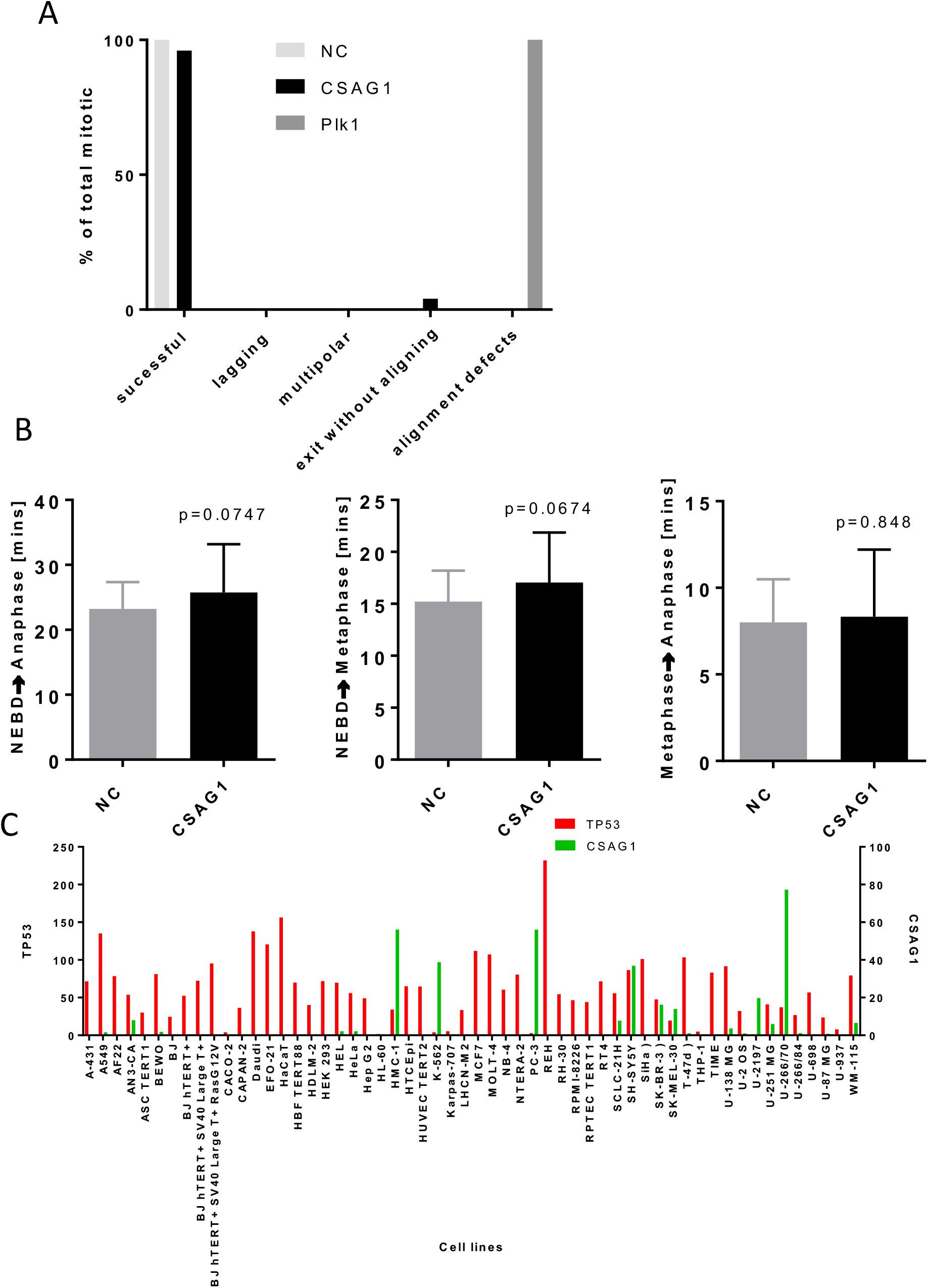
RPE1 cells are resistant to CSAG1 depletion phenotypes. **(A)** Mitotic phenotypes were examined in RPE1 cells treated with NC siRNA, siRNA to CSAG1, or siRNA to Plk1. **(B)** Elapsed times from NEBD-metaphase, metaphase to anaphase were determined in cells from A. Totals of >60 cells per siRNA treatment were examined. The Mann-Whitney test was used for statistical analysis. RPE1 cells do not show any discernable mitotic phenotype after CSAG1 depletion beyond a marginal increase in mitotic duration. (**C**) Numbers of p53 and CSAG1 transcripts in different cell lines were plotted from publicly available data (www.sciencemag.org/content/356/6340/eaal3321/suppl/DC1). No correlation between p53 and CSAG1 transcript levels was evident.

**Supplemental video 1:** HeLa cells stably expressing GFP-Tubulin were transfected with siRNA targeting the UTR of CSAG1. Cells were labeled with sirDNA dye to visualize DNA and imaged every 7 minutes. Video shows a HeLa cell from NEBD to mitotic exit. Spindle pole fragments causing metaphase plate to bend.

